# Force profile of the two-handed hardstyle kettlebell swing in novice older adults: an exploratory profile

**DOI:** 10.1101/2021.05.17.444430

**Authors:** Neil J. Meigh, Wayne A. Hing, Ben Schram, Justin W.L. Keogh

**Author notes:** Corresponding Author: Neil Meigh, Bond Institute of Health & Sport, 2 Promethean Way, Robina, QLD, 4226, Australia.

## Abstract

**Background:** Understanding the force profile of an exercise increases clinical confidence when assessing the benefits and potential risks of a prescribed exercise. This exploratory study presents the force profile of the hardstyle kettlebell swing in novice older adults and compares peak force with kettlebell deadlifts. These data will help inform healthcare providers and coaches who are considering prescribing kettlebell exercises for older adults.

**Methods:** Thirty-five community-dwelling males and females (59-79 years) were recruited, from applicants to participate in the BELL trial. Two-handed hardstyle swings were performed with 8-16 kg. Deadlifts were performed with 8-24 kg and 8-32 kg for females and males, respectively. Ground reaction force was obtained from a floor-mounted force platform. Pairwise comparisons of peak force, forward force, rate of force development, swing cadence, sex, and kettlebell mass, were investigated for the kettlebell swing, with representative force-time curves described. Pairwise comparisons of peak force, sex and kettlebell mass were investigated for the deadlift, with comparisons of peak force, kettlebell mass, and sex, between swings and deadlifts.

**Results:** For kettlebells up to 16 kg, paired samples T-tests show a large exercise effect (δ > 1.4) with peak force higher for swings than deadlifts. Data shows: (i) higher peak force during swings than deadlifts (δ = 1.77), reaching 4.5 (1.0) N.kg^-1^, (ii) peak force during an 8 kg swing was greater than a 32kg deadlift, (iii) negligible difference in normalised peak force between males and females performing kettlebell swings, but a moderately large effect size during deadlifts (males > females, δ = 0.69), (iv) mean rate of force development of 19.9 (4.7) N.s^-1^.kg^-1^ with a very weak, positive correlation with kettlebell mass (*y* = 14.4 + 0.32*x*), and trivial effect of sex, (v) mean forward force equal to 5.5% of vertical force during swings, increasing from 3.8 (1.6) % with 8 kg to 7.1 (2.6) % with 16 kg.

**Conclusion:** During kettlebell swings, there is negligible difference in normalised net peak force between novice males and females using the same absolute loads. Where ground reaction force is a therapeutic target, kettlebell swings with an 8 kg kettlebell could have similar effects to much heavier deadlifts (>24 kg). Kettlebell swings performed with lighter loads, could provider similar therapeutic value to much heavier deadlifts, and may be a more appealing, affordable, and convenient option for older adults.

## Introduction

To date, very few clinical trials have been conducted to investigate the effects of kettlebell training in older adults. The limited results however, have been encouraging, with significant improvements in strength, muscle mass, and peak expiratory flow, in older females with sarcopenia ^(1)^, and improvements in functional performance in older adults with Parkinson’s disease ^(2)^. Small and portable, kettlebells have been recommended for their ease of use, affordability, and for being less intimidating than other equipment ^(3)^. These are attractive features for clinicians and researchers who have used kettlebells in rehabilitative programs for older adults ^(4)^.

An important component of therapeutic exercise is the choice of exercise and load, and knowledge of resultant force, which can be used as a quasi-measure of the force applied to target tissues. A successful clinical outcome of therapeutic exercise may depend upon the correct manipulation of force to stimulate a desired adaptative response ^(5)^. In older adults, repetitive loading is used in the management of hip and knee arthritis, back pain, and osteoporosis ^(6, 7)^. Healthcare providers must first do no harm. Although the risk of adverse events during exercise appear to be low ^(8)^, providers must remain mindful of inappropriate volume, intensity, or load, which could result in acute injury, excessive muscle soreness, or irritate an already painful condition such as osteoarthritis. Understanding the force profile of an exercise can be a helpful, often necessary starting point, when prescribing exercise programs for rehabilitation and performance.

For an older population, ground reaction force (GRF) and rate of force development (RFD) are important measures of functional decline ^(9, 10)^. Ground reaction force is associated with strength and power at the hip and knee ^(11)^, gait speed ^(10)^, falls risk ^(12)^, and may be a better predictor of functional decline than both grip strength ^(9)^ and the five-times Sit-to-Stand test ^(11)^. Age-related loss of muscle power is associated with an increased incidence of falls ^(13, 14)^, but this can be improved with explosive resistance training ^(15, 16)^. Movement patterns observed in a Sit-To-Stand, floor transfer, and reaching overhead, are also seen in kettlebell training; the most widely recognised type being the ‘hardstyle’ swing introduced by Pavel Tsatsouline ^(17)^.

The hardstyle swing is characterised by i) ballistic movement of the lower limb, ii) a ‘hip-hinge’ motion, iii) four distinct phases, and iv) the kettlebell reaching chest-height at mid-swing. The start of a swing cycle begins with the subject in a universal athletic position, mid-forearms resting on the top of the thighs, and the kettlebell located between the legs, posterior to the hips. The kettlebell follows an upward arc of trajectory, accelerated by explosive extension of the hips and knees (phase 1). The second phase, called ‘float’, involves only passive flexion of the shoulders, which occurs between full extension of the lower limbs and the kettlebell reaching its maximum vertical displacement (mid-swing). Phase three, called ‘drop’ is the reverse of phase 2 - passive shoulder extension to commencement of hip flexion. The fourth and final phase is ‘braking’, in which the lower limbs decelerate the kettlebell back to the start position ^(18)^. A hardstyle swing is believed to improve lower limb power, and by association, the ability to apply GRF. Therefore, the kettlebell swing might positively influence lower limb function in older adults, reduce risk of falls, and be an ideal tool to promote healthy ageing ^(19)^.

A recent review ^(19)^ identified no standards or guidelines to inform healthcare providers of how to use kettlebells safely and confidently with older adults. Additionally, current resistance training guidelines for older adults ^(20-22)^ do not translate to hardstyle kettlebell practices as a single exercise ‘set’ may involve 100 repetitions and last several minutes. Selection of a kettlebell is not typically based on a percentage of ‘1 repetition maximum’, and simply adding a kettlebell to an exercise ^(4)^ is not representative of ‘kettlebell training’. It is a safety concern for providers wanting to prescribe a ballistic, free weight resistance training program for older adults, without guidelines and for which the force profile, acute responses, and risks are unknown.

Hardstyle literature suggests that outcomes, such as improvements in hip extension power, are dependent upon technique ^(17, 23)^. Kinematic differences between novice and expert have been described ^(24)^ with significant, large effect size differences in GRF between novice and instructor ^(18)^, however, these data are from healthy young adults. Data from young active populations are informative and provide important reference points ^(25-30)^, but these cannot be generalised to older adults, especially those who insufficiently active, or living with chronic health conditions. Our research question centers on ‘force’, due to its emphasis and influence in kettlebell practice - “An experienced girevik is playing a game of “force pool,” expertly rebounding the force generated by the clean from the ground.” ^(17)^, its’ role in mechanotransduction, and informing optimal dosing strategies for therapeutic exercise prescription ^(31, 32)^.

For adults under 65 years of age, insufficient physical activity is defined as not completing 150 minutes of moderate to vigorous physical active across five or more days in the last week, which includes muscle strengthening activities twice a week. For adults 65 years and over, it is 30 minutes of physical activity per day on five or more days in the last week, incorporating muscle strengthening activities ^(33)^. A profile of the kettlebell swing in older adults is therefore warranted.

The aim of the study is to report the force profile of the two-handed hardstyle kettlebell swing in novice older adults. Peak ground reaction force, RFD, forward force, and swing cadence would be compared by sex, with force-time curves (FTCs) reported and described. Peak GRF during swings would be compared with GRF during kettlebell deadlifts, with findings used to inform the BELL trial intervention (www.anzctr.org.au ACTRN12619001177145).

## Materials & Methods

### Participants

A total of 17 males and 18 females aged 58-79 years, who had volunteered to participate in the BELL trial, were recruited. Thirty-two participants identified as insufficiently active ^(33)^ and had not engaged in a structured exercise program for at least nine months. None of the participants had previously used kettlebells. Participants were free from injury and did not disclose any health or medical conditions considered to be a high risk ^(34)^. After a thorough explanation of the study aims, protocols, and potential risks, participants provided written informed consent. A copy of the explanatory statement and informed consent forms are included in Appendix 6 and 7. Ethical approval for this study was granted by Bond University Human Research Ethics Committee (NM03279).

### Protocol

Data were collected from the University biomechanics laboratory, each participant attending two, 1-hour sessions, on consecutive weeks. Two-handed kettlebell swings to chest-height were performed on a floor-mounted force flatform (AMTI, Watertown, NY, USA) recording GRF at 1000 Hz using NetForce software (AMTI, USA). Participant body mass was captured by the force plate from a period of quiet standing. Tri-plantar force variables were obtained from the floor-mounted force platform. The variables of interest were peak GRF, forward force, dynamic RFD, swing cadence, sex, and kettlebell mass. Participants performed a single set of 12 repetitions with each kettlebell, with the middle 10 repetitions used for analysis. The first and last swings in each set, involving the dead-start/dead-stop position, were excluded from analysis. A custom program (Microsoft Excel, Version 2012) was used to calculate peak force during each swing cycle of the set, with values manually assessed and verified against the corresponding FTC. To obtain net peak force, system weight (body mass + kettlebell mass multiplied by gravity) was subtracted from the square root of squared and summed data:

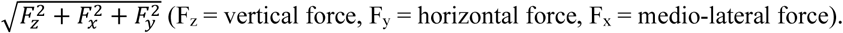

The swing pattern is reported as previously described ^(18)^. The back or bottom position of the swing was used as the start of each swing cycle. Dynamic RFD (N.s^-1^) during hip extension (propulsion) was calculated as the change in resultant GRF during this first phase, divided by elapsed time. This was normalised to body mass (N. s^-1^.kg^-1^) and reported as the mean of 10 swings. Cadence, in swings per minute (SPM), was calculated from the average time between the peak force of hip extension in each swing cycle – 60s / (Time_peak10_ - Time_peak1_ / 9). Peak force was reported as resultant force unless stated otherwise.

### Procedure

During the first session, participants received instruction on how to perform the hardstyle swing as previously described ^(18, 35)^. The following drills served as a warm-up; (1) unloaded hip-hinge - standing at a distance roughly equal to the length of their foot away from a wall, participants used the wall as an external focus of attention (target for their buttocks to touch) to practice the ‘hinge’ movement, before returning to an upright position. (2) performing the 4 phases of a swing cycle ^(18)^ without a kettlebell, led by the instructor to a 4-count (propulsion, float, drop, braking). (3) double knee extension swings ^(36)^ with 8 kg, to familiarise themselves with the external load, movement of the kettlebell through the legs, and weight shift. (4) towel-swings ^(17)^ with 4-8 kg, to differentiate a ‘swing’ from a ‘lift’, and (5) two to three sets of instructor-led swings performed at cadence of approximately 40 SPM, to illustrate the intended ballistic nature of the hardstyle swing.

Force USA competition kettlebells from 8-32 kg of standardised dimensions were used, increasing in 2 kg increments to 24 kg, then 4 kg increments to 32 kg. Target loads for swings and deadlifts were pre-selected based on the lead investigator’s experience as a kettlebell instructor; 12 kg and 24 kg for females, and 16 kg and 32 kg for males, respectively. Sets of swings began and ended in a ‘dead-start’ position (Fig. 1 A). Deadlifts were performed with feet positioned either side of the kettlebell (Fig. 1 D) as previously described ^(37)^. Only peak resultant force was analysed for deadlifts.

**Figure 1.**
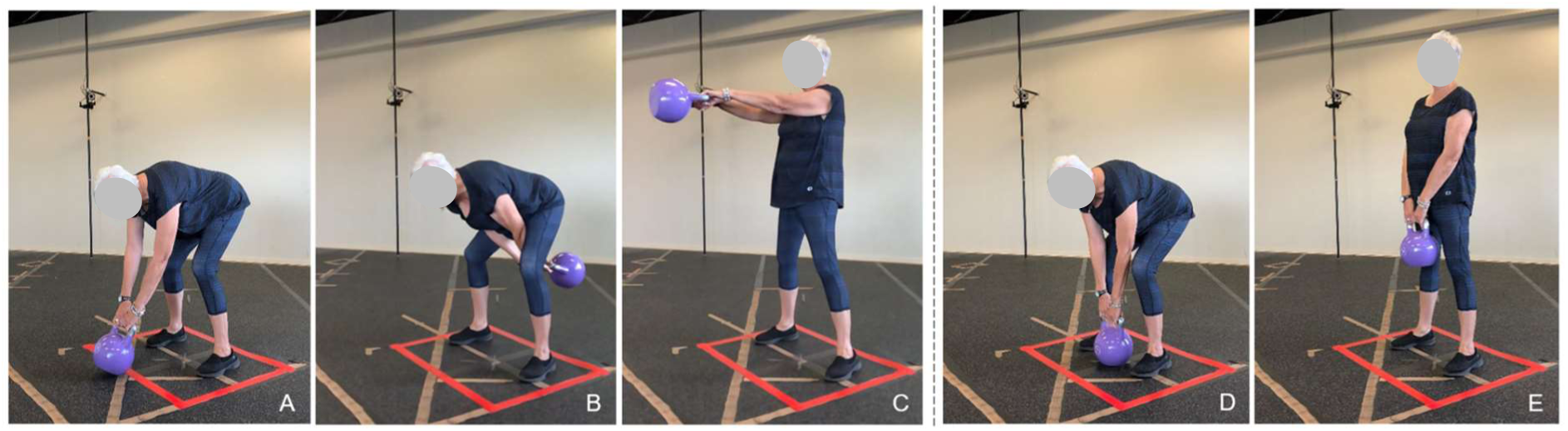
Key positions during the two-handed hardstyle kettlebell swing and kettlebell deadlift. (A) dead-start / dead-stop for a swing, (B) start and end position of single swing cycle, (C) mid-swing, (D) dead-start for a kettlebell deadlift, (E) top position of the deadlift. Red tape marks the boundary of the force platform.

A target range of kettlebells was set for both exercises, however, the physical capacity of the participants was unknown. A maximum of 16 kg was set for swings, and 32 kg for deadlifts. It was anticipated that some individuals may be unable (or unprepared) to use all the kettlebells or, may wish to go beyond the target range. Females were permitted to perform swings with 14 kg and 16 kg if they requested to do so, and had confidently performed swings using the lighter kettlebell with acceptable technique. As an exploratory investigation, standardising kettlebells across all participants was not required, and additional data about physical capacity was beneficial to the larger clinical trial within which this study was subsumed.

### Statistical analyses

Descriptive statistics were reported for normally distributed continuous variables. Measures of centrality and dispersion are presented as mean (SD) and verified for normality using normal Q-Q plots, histograms, and Shapiro-Wilk tests. Differences between males and females, swings, and deadlifts, were analysed using independent samples T-tests and paired samples T-tests, respectively, subject to the normality assumptions being met. For these analyses, mean Difference (MD) is presented with 95% confidence interval (CI). An exploratory data analysis of pairwise comparisons, involving exercise (swings vs deadlift), sex (male vs female) and kettlebells (8-32kg), resulted in 60 pairwise comparisons. A Bonferroni correction for multiplicity, provided an adjusted *p*-value of 0.0008 as the threshold for statistical significance. Hypothesis confirmation cannot be performed with exploratory data. Due to the exploratory nature of the study, not all conditions were met by all participants, giving rise to incomplete cases; thus, although the repeated measures ANOVA and other variations like the mixed model ANOVA were considered, they were not carried out as these methods would have discarded incomplete cases. Effect sizes were calculated and interpreted using Lenhard and Lenhard ^(38)^ and Magnusson ^(39)^, quantified as trivial, small, moderate, large, very large, and extremely large where effect size < 0.20, 0.20 - 0.59, 0.60 - 1.19, 1.20 - 1.99, 2.0 - 3.99 and ^3^ 4.0 respectively ^(40)^. Probability of superiority has been used to illustrate the Cohen’s d effect size (δ), representing the chance that a person from group A will have a higher score than a person picked at random from group B ^(39)^. Linear regression was used to calculate the regression coefficients between the independent variable load, and dependent variables net peak force, and rate of force development. Correlations were investigated using Pearson product-moment correlation coefficient, with preliminary analyses performed to ensure no violation of the assumptions of normality, linearity, or homoscedasticity. Statistical analyses were performed using SPSS (version 26.0; SPSS Inc., Chicago, IL, USA), and *p* < 0.05 was used to indicate statistical significance. Readers are reminded of the limited utility of exploratory *p*-values in data exploration for hypothesis generation ^(41)^, thus directed to the effect size of reported pairwise comparisons.

## Results

Participant demographics are presented in Table 1. All participants performed swings with 8-12 kg. Nine females performed swings with 14 kg, and two females performed swings with 16 kg. Sixteen males performed swings with 14 kg, and 15 males performed swings with 16 kg.

**Table 1.** Participant characteristics

| Characteristic | Older Males | Older Females |
| --- | --- | --- |
| Number | 17 | 18 |
| Age (years) | 68.5 (4.2) | 68.1 (4.0) |
| Height (cm) | 176.8 (7.8) | 163.9 (5.4) |
| Weight (kg) | 90.2 (14.9) | 71.7 (11.5) |
| BMI (kg/m<sup>2</sup>) | 29.1 (2.5) | 26.6 (4.5) |

*Data are presented as mean (SD), unless otherwise specified.*

### Net peak ground reaction force

There was a large (δ = 1.77) difference in peak force between the kettlebell swing and kettlebell deadlift (Fig. 2 and Table 2), with the mean probability of superiority for the kettlebell swing across loads of 8-16 kg being 89.5%. The relationship between net peak force and kettlebell mass was investigated. In both cases, there was a very weak (*r* < 0.30) positive correlation between the two variables. Regression coefficients to predict net peak force, where *x* = kettlebell mass in kg.

**Table 2.** Net peak ground reaction force for swings and deadlifts (N.kg^-1^)

| KB weight (kg) | Mean (SD) |  | MD (95% CI) | *t* | *p* | Cohen’s |  |
| --- | --- | --- | --- | --- | --- | --- | --- |
| | Swing | Deadlift | | | | $\delta$ | Probability of superiority (%) |
| 8 | 3.9 (1.0) | 2.8 (0.6) | 1.2 (0.8, 1.6) | 29 (6.110) | <0.001 | 1.42 | 84.2 |
| 10 | 4.3 (1.0) | 2.8 (0.7) | 1.5 (1.2, 1.9) | 29 (9.408) | <0.001 | 1.76 | 89.3 |
| 12 | 4.4 (0.8) | 2.8 (0.6) | 1.5 (1.2, 1.9) | 29 (9.927) | <0.001 | 1.99 | 92.0 |
| 14 | 4.7 (1.0) | 3.0 (0.6) | 1.7 (1.3, 2.1) | 22 (8.887) | <0.001 | 2.00 | 92.1 |
| 16 | 4.5 (1.0) | 3.1 (0.6) | 1.4 (0.8, 2.0) | 15 (5.067) | <0.001 | 1.68 | 88.3 |

**Figure 2.**
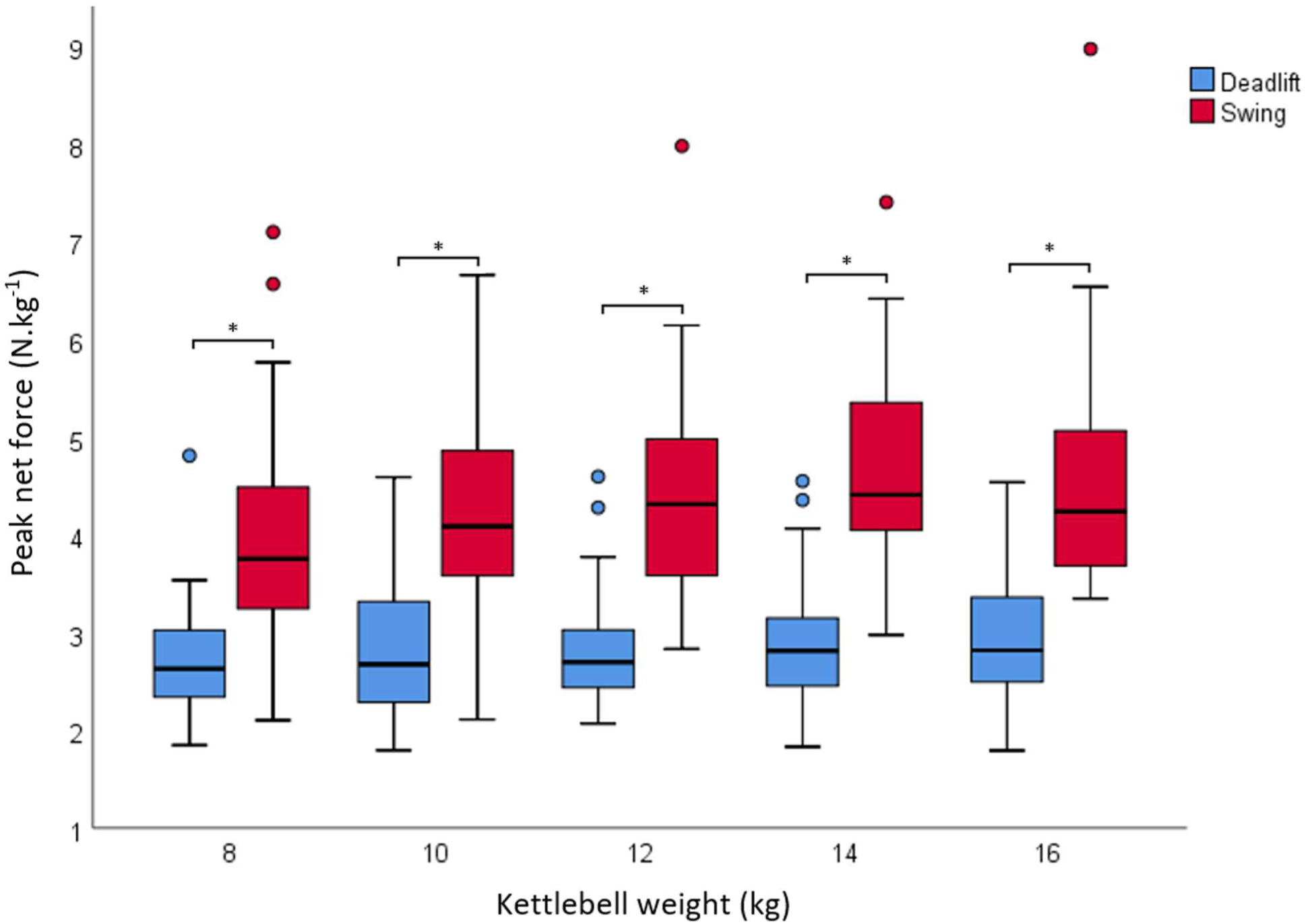
Net peak ground reaction force between kettlebell swings and deadlifts. Box-and-whisker plot of large difference in net peak ground reaction force between kettlebell swings and deadlifts. Box represents median observations (horizontal rule) with 25th and 75th percentiles of observed data (top of bottom of box). Length of each whisker is 1.5 times interquartile range. Data points that are more than 1.5× interquartile range are represented by “o” (Tukey’s outlier detection method). * = *p* <0.05

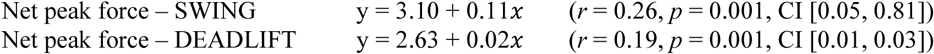

Only trivial-to-small effects were found between males and females performing kettlebell swings up to 16 kg, however sex influenced peak force during kettlebell deadlifts. From 8-24 kg, peak force during deadlifts was moderately higher (δ = 0.69) for males than females, with a 68.7% probability of superiority. There was a difference between males and female using 8, 12 and 14 kg kettlebells, with the mean probability of superiority being 71.2% (δ = 0.79) (Fig. 3 and Table 3).

**Table 3.** Net peak ground reaction force for males and females during kettlebell deadlifts (N.kg^-1^)

| KB weight (kg) | Mean (SD) |  | MD [95% CI] | *t* | *p* | Cohen's |  |
| --- | --- | --- | --- | --- | --- | --- | --- |
| | Male | Female | | | | $\delta$ | Probability of superiority (%) |
| 8 | 3.0 (0.7) | 2.5 (0.4) | 0.4 [<0.05, 0.87] | 28 (2.112) | 0.044 | 0.77 | 70.7 |
| 10 | 3.0 (0.7) | 2.6 (0.6) | 0.3 [-0.2, 0.8] | 28 (1.318) | 0.198 | 0.48 | 63.3 |
| 12 | 3.1 (0.7) | 2.6 (0.4) | 0.5 [<0.05, 0.9] | 28 (2.242) | 0.033 | 0.82 | 71.9 |
| 14 | 3.1 (0.6) | 2.7 (0.6) | 0.4 [<0.05, 0.9] | 28 (1.970) | 0.059 | 0.72 | 69.5 |
| 16 | 3.2 (0.6) | 2.7 (0.6) | 0.5 [<0.05, 0.9] | 28 (2.114) | 0.044 | 0.77 | 70.7 |
| 18 | 3.1 (0.6) | 2.8 (0.6) | 0.3 [-0.1, 0.7] | 28 (1.417) | 0.168 | 0.52 | 64.3 |
| 20 | 3.2 (0.6) | 2.8 (0.6) | 0.4 [<0.05, 0.8] | 28 (1.899) | 0.068 | 0.69 | 68.7 |
| 22 | 3.2 (0.6) | 2.8 (0.5) | 0.3 [<0.05, 0.7] | 28 (1.797) | 0.083 | 0.66 | 68.0 |
| 24 | 3.3 (0.6) | 2.9 (0.6) | 0.4 [<0.05, 0.8] | 28 (1.861) | 0.073 | 0.77 | 10.7 |
| 28 | 3.4 (0.7) | 2.7 (0.2) | 0.8 [-0.3, 1.8] | 14 (1.537) | 0.147 | 1.16 | 79.4 |
| 32 | 3.6 (1.0) | 2.6 (0.5) | 1.0 [-0.5, 2.5] | 12 (1.408) | 0.184 | 1.08 | 77.7 |

**Figure 3.**
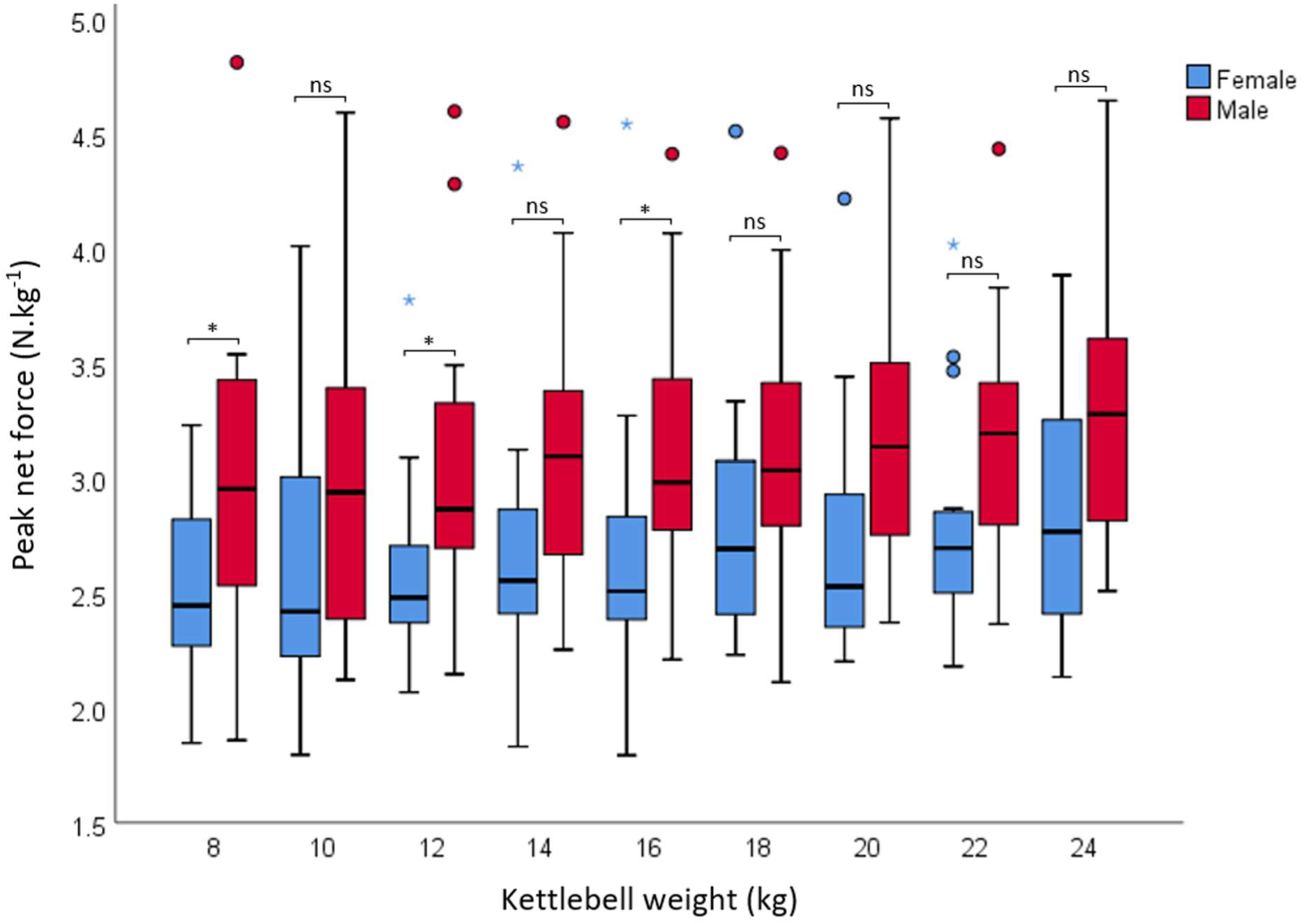
Effect of sex on peak force during kettlebell deadlifts. Moderately large effect of sex on peak force during kettlebell deadlifts with 68.7% mean probability of superiority (males > females) with 8-24 kg kettlebells. Data points that are more than 3× interquartile range are represented by “☆”. * = *p* <0.05, ns = *p* >0.05

### Rate of force development

Rate of force development was not significantly different between consecutive kettlebells. There was a difference (δ = 0.5 - 0.7) in RFD between the 8 kg and 14 kg kettlebell, and the 8kg and 16 kg kettlebell: MD = 2.2 (0.5, 3.9), *t* = (24) 2.721, *p* = 0.012, and MD = 3.89 (1.59, 6.20), *t* = (16) 3.577, *p* = 0.003, respectively (Fig. 4 and Table 4).

**Table 4.** Rate of force development during kettlebell swings for males and females (N.s^-1^.kg^-1^)

| KB weight (kg) | Mean (SD) |  | MD [95% CI] | *t* | *p* | Cohen's |  |
| --- | --- | --- | --- | --- | --- | --- | --- |
| | Males | Females | | | | $\delta$ | Probability of superiority (%) |
| 8 | 15.5 (4.9) | 18.7 (3.6) | 3.3 [0.3, 6.6] | (33) 2.255 | 0.031 | 0.75 | 70.2 |
| 10 | 16.7 (4.7) | 17.8 (3.7) | 1.1 [-1.8, 4.0] | (33) 0.776 | 0.443 | 0.26 | 57.3 |
| 12 | 18.3 (6.5) | 18.1 (3.2) | -0.2 [-3.7, 3.3] | (33) -0.111 | 0.913 | 0.04 | 51.1 |
| 14 | 18.4 (4.7) | 18.8 (4.1) | 0.5 [-3.4, 4.3] | (23) 2.47 | 0.807 | 0.09 | 52.5 |
| 16 | 19.3 (5.2) | 21.8 (9.3) | 2.5 [-6.5, 11.5] | (15) 0.560 | 0.560 | 0.45 | 62.5 |

**Figure 4.**
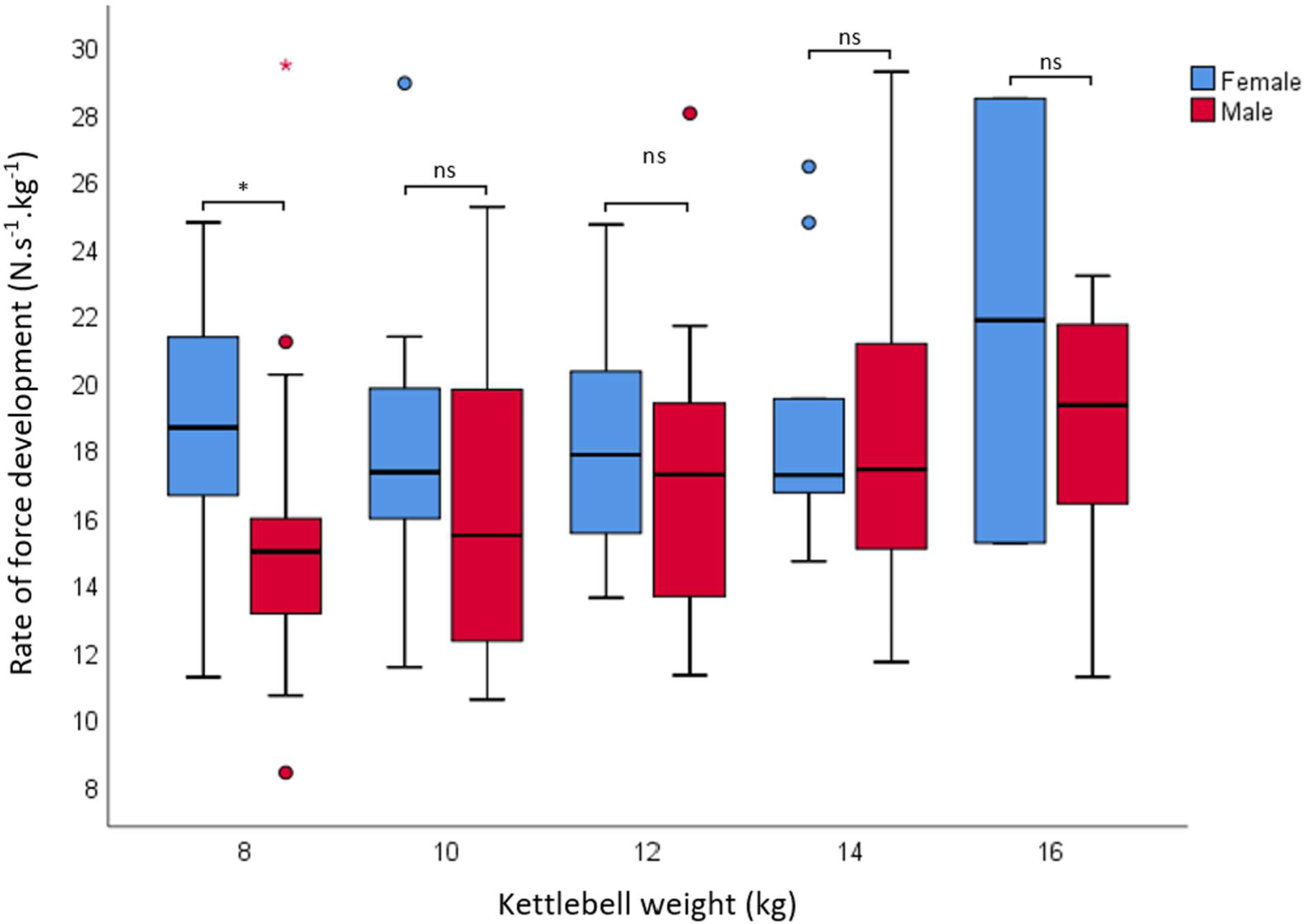
Effect of sex on RFD. Trivial effect of sex on RFD. * = *p* <0.05, ns = *p* >0.05. One outlier at 16 kg and one extreme outlier at 12 kg not shown.

Analysis of the effect of sex on RFD revealed a difference between males and females, but only with 8 kg (Fig. 4). Except for the 2 females who swung 16 kg, paired samples analysis revealed no significant increase in RFD between the lightest and heaviest kettlebells for males or females. The relationship between RFD and kettlebell mass was investigated. Linear regression shows a very weak, positive correlation between the two variables where:

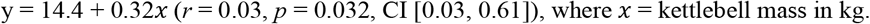

### Forward force

Mean forward force was 0.055 (5.5%) of the vertical GRF during the kettlebell swings. There was a small (δ = 0.34) increase in the ratio of forward to vertical GRF between 8 and 16 kg, MD = 0.33 (0.21, 0.45), *t* = (14) 5.890, *p* = <0.001. The trivial to small differences between males and females were not significant at any kettlebell (Fig. 5). Between 8 and 14 kg, the absolute change in ratio of horizontal to vertical force was 2.5% for females, increasing from 3.7% to 6.9%. For males, the absolute change was 2.6%, from 3.8% with 8 kg to 7.3% with 16 kg.

**Figure 5.**
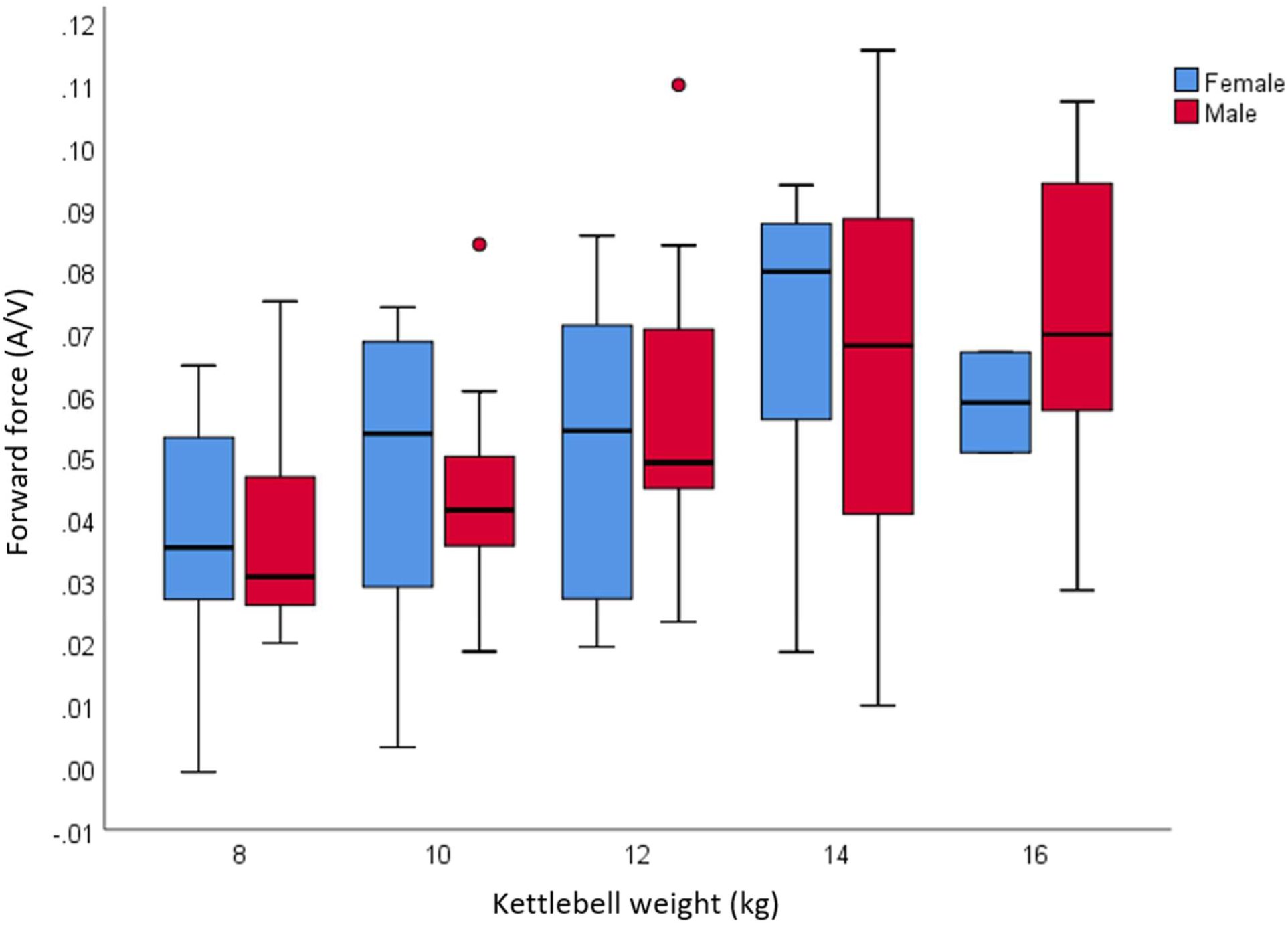
Effect of sex on forward force. No significant effect of sex on small increases in forward force with increasing kettlebell mass. A/V = ratio of *anterior* (forward) horizontal to *vertical* GRF.

### Cadence

Mean swing cadence up to 16 kg for the kettlebell swings was 35.7 (2.3) SPM, range = 29.2 to 41.2. There was a difference (*p* < 0.05) in cadence between 10 and 12 kg and 12 and 14 kg swings, however, means remained between 35 and 36 SPM. The effect of sex on cadence revealed differences between males and females with all kettlebells (Fig. 6). Between 8 and 16 kg, there was a moderately large (δ = 0.9) effect size, with a mean difference of 1.9 SPM and 73.8% probability of superiority (females > males).

**Figure 6.**
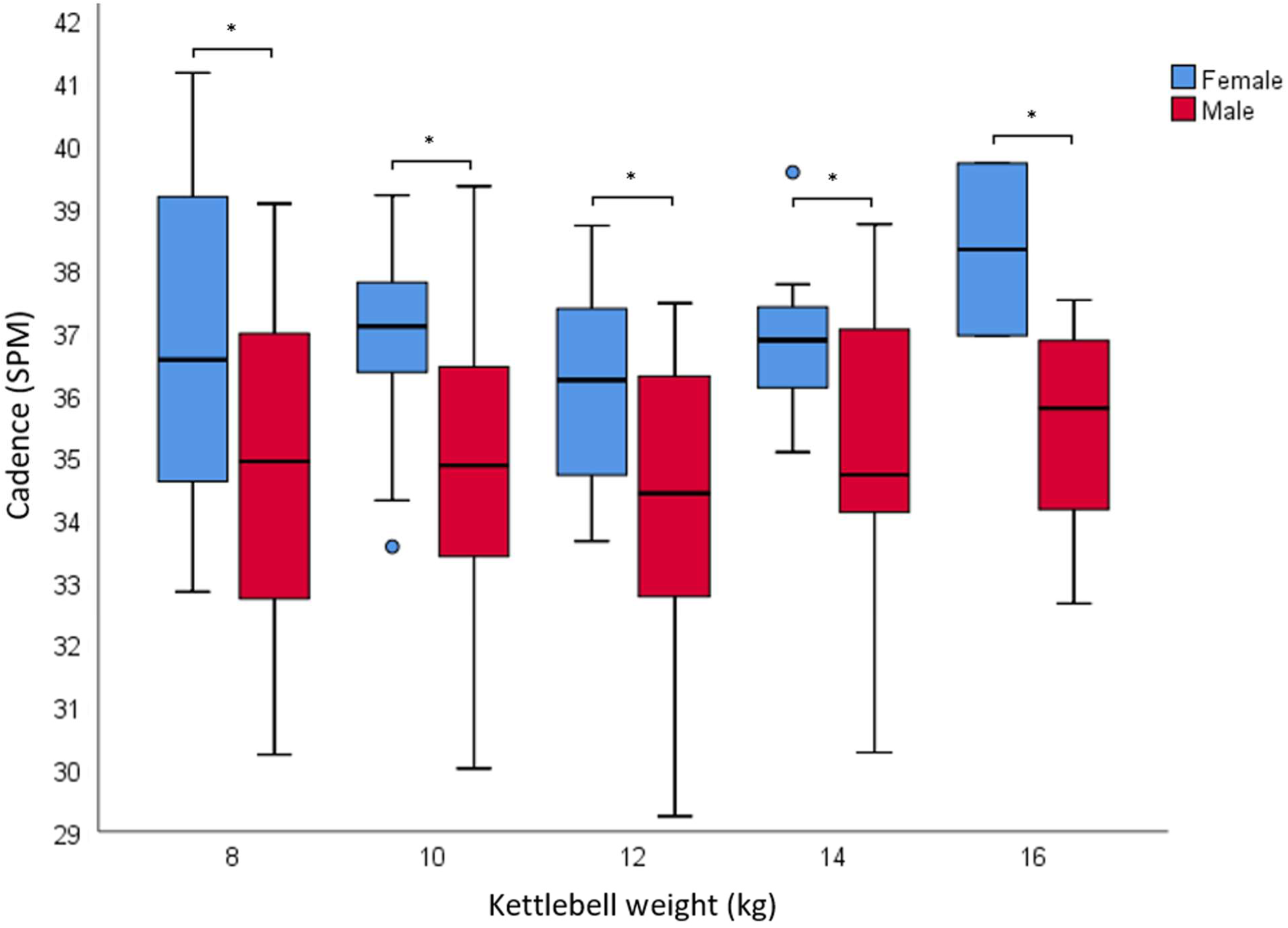
Effect of sex on swing cadence. Moderately large effect of sex on swing cadence. SPM = swings per minute. * = *p* <0.05

### Force-time curve characteristics

Three distinct movement strategies emerged, which were inconsistent with a proficient swing ^(18)^. 1) lifting the kettlebell with the upper limbs, 2) a whole-body effort to perform the movement, 3) controlling the kettlebell’s decent from mid-swing, with little visual dissociation between upper and lower limbs, in which the body appeared to collapse downward with the kettlebell’s vertical displacement. Representative FTCs recorded from two participants which illustrate these movement strategies, are shown in Fig. 7. A double-peak (Fig. 7 A) is consistent with a sequential movement, initially involving explosive hip extension, immediately followed by active shoulder flexion to lift the kettlebell to the desired height. A wider, multi-peaked profile (Fig. 7 B) is consistent with a more simultaneous movement, and lack of ballistic hip extension during propulsion. Absence of a second smaller force peak (shown in Fig. 7 A) is consistent with not allowing the kettlebell to drop from mid-swing. These profiles illustrate the effect of technique on force and cadence. The ballistic pattern more consistent with a proficient hardstyle swing (A), resulted in higher net peak force (368.2 N vs 257.1 N), higher force as a percentage of bodyweight (47.5 % vs 33.8 %), a larger proportion of horizontal forward force (6.8% vs 2.9 %), and faster swing cadence (36.4 SPM vs 34.1 SPM).

**Figure 7.**
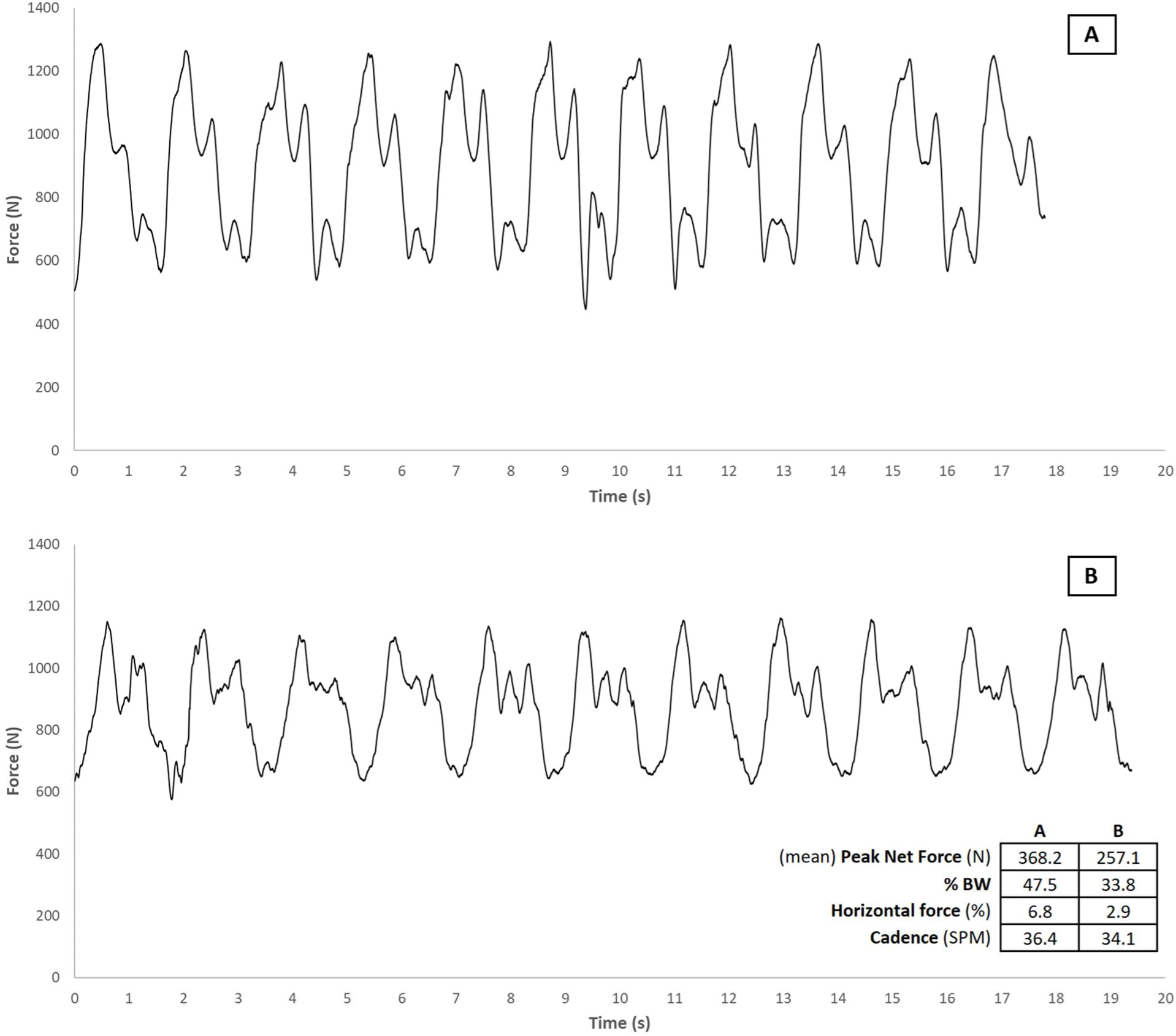
Representative force-time curves from two distinct movement strategies used by novice older adults to perform a two-handed hardstyle kettlebell swing.

## Discussion

Knowledge of GRF and other kinetic variables enables healthcare providers to make informed decisions about the potential benefits and risks of an exercise. A properly designed resistance training program for older adults, includes multi-joint exercises such as the kettlebell swing and kettlebell deadlift, is individualised, periodised, progressive, and includes appropriate technique instruction ^(22)^. A review ^(19)^ was unable to identify descriptive data of kettlebell exercises from older adults for any of these variables. Results of the present study provide healthcare providers with additional data, to make more informed choices about the potential therapeutic use of kettlebell swings for older adults.

### Ground reaction force

It was unsurprising that net peak GRF was higher for swings than deadlifts. Force is a product of mass and acceleration and increases with speed in all environmental conditions ^(42)^. Net peak ground reaction was 5% higher during swings with an 8 kg kettlebell than deadlifts with 32 kg, and 35% higher than deadlifts with 16 kg. If GRF is the desired therapeutic stimulus, an 8 kg kettlebell swing could be a better option than a heavier deadlift. Other factors such as reduced cost, ease of storage and transport, and availability in a variety of general-purpose stores, would likely make kettlebell swings with an 8 kg kettlebell, a less intimidating and more attractive option than deadlifts, which would typically be performed with much heavier kettlebells.

Data from proficient hardstyle swings performed by an Instructor, show ground reaction force can exceed 2× body mass with an 8 kg kettlebell ^(18)^. For the novice older adults in the present study, net peak force during swings with an 8kg kettlebell was 3.9 (1.0) N.kg^-1^, compared to 8.85 N.kg^-1^ for the younger instructor previously reported.

With appropriate training, it is feasible that ground reaction force in novice older adults could be increased from 4 N.kg^-1^ to 8 N.kg^-1^ during hardstyle swings with 8 kg. This highlights a potential therapeutic opportunity in an older population, and possible goals for exercise prescription and measurement.

Lower limb strengthening, for example, is recommend in the management of a host of chronic diseases, including symptomatic knee osteoarthritis ^(43, 44)^. During the kettlebell swing, knee flexion is limited ^(24)^ with no impact force - mechanisms commonly reported to irritate symptomatic arthritic knees. At a cadence of 40 swings per minute, a training load volume of 1,600 kg can be achieved in as little as five minutes with an 8 kg kettlebell. Safe, easily achievable, and readily scalable, the swing could be an effective and efficient option where time, motivation, or desire, might be limited. Ground reaction force relative to bodyweight during an 8 kg swing is comparable to a bodyweight squat performed at a medium to fast pace ^(42)^, but the number of bodyweight squats a healthcare provider might prescribe for an older adult, is likely to be far less than the number of swings that could be performed in the same time period. This may have implications for the prescription of training load volume.

More than half of the female participants voluntarily swung a kettlebell heavier than the intended 12 kg maximum. It was also evident that some of the males had capacity to swing kettlebells heavier than 16 kg. As these participants were all novices with respect to kettlebell training, this was very encouraging to the potential feasibility of kettlebell swings in this population. Whether there is any physical benefit from using heavier kettlebells has yet to be determined, particularly as effort increases disproportionately to kettlebell mass, and change in GRF is small ^(18)^. If the clinical benefits can be gained with a lighter rather than heavier kettlebell, it appears the potential risks of swinging a heavier kettlebell could quickly outweigh the reward. To minimise the risk of injury, it seems reasonable that 12-16 kg would be “sufficient” for most older adults, with other program variables available to alter the physiological demand, such as the work:rest ratio, altering technique and cadence, and option to use one-arm instead of two.

Males have a higher volume of lean muscle mass than females ^(45-47)^, so it was anticipated that peak force would be higher for males when performing both kettlebell exercises. This was true for deadlifts but not for swings, which was surprising. There was a consistent moderate effect size with an almost 70% mean probability of superiority of peak force being higher for males. There was a sex effect for deadlifts, however the lack of such significance for swings is currently unclear.

There was a very large mean effect size (δ = 3.06) in peak force between the older adults in this study and data reported from an instructor ^(18)^. For older adults, the average peak force during swings was almost half that of the instructor (52.1%), with a mean difference of 4.2 N.kg^-1^, highlighting scope for potential improvement from training. Previous research has shown no significant difference in peak GRF, during lifting and standing tasks, between younger and older adults ^(48, 49)^, but moderate differences between older community-dwelling males and females ^(11)^. In the absence of sex-specific health or medical conditions, these results suggest that choice of kettlebell should be influenced by bodyweight, strength and/or stature, rather than sex.

Novice starting kettlebell loads for the kettlebell swing are commonly based upon recommendations which are not specific to the swing or older adults ^(17)^. Researchers have largely followed these recommendations, however, some investigators have used kettlebells as low as 3 kg ^(50-54)^, heavier kettlebells up to 48 kg ^(30, 36, 55-60)^, or a percentage of bodyweight e.g., 10-20% ^(61, 62)^. Results from this study suggest that healthy novice females, regardless of age and training status, with some supervision and guidance, may be able to perform swings safely a with 12-14 kg kettlebell, if they feel comfortable doing so.

### Forward force

“Street wisdom” ^(25)^ suggests that most of the applied force is in the forward horizontal direction. This belief has influenced training practices, which include visualising the kettlebell being projected forward during the propulsion phase, and forcibly snapping the knees into full extension. A kinetic analysis of a proficient hardstyle swing, revealed horizontal forward force to be approximately 10-15% of the magnitude of vertical force, with kettlebells from 8 to 32 kg ^(18)^. One of the most striking findings from the present study, was that only around 5% of the ground reaction of a swing was directly horizontally forward. Mean forward force increased from just 3.8% with 8 kg to 7.1% with 16 kg. A small proportion of forward force was expected, however, a mean of 5.5% with few swings exceeding 10% was much less than previous research in younger adults ^(27)^. This finding may be due to age-related decline in dynamic balance.

There was a very large (δ = 2.05) average effect size in the magnitude of forward force between older adults and instructor ^(18)^. Although the absolute mean difference was less than 5%, the relative difference was just over half (55.1%). In some cases, horizontal force at peak ground reaction was zero, and in a few instances, negative, albeit negligible in magnitude i.e., indicating that the kettlebell was in effect being pulled backward rather than projected forward at the point of peak ground reaction. This is an interesting biomechanical scenario, given the kettlebell’s anteroposterior arc of motion, and street wisdom regarding forward force. Tsatsouline describes the motion of the lower limbs during a kettlebell swing as akin to a standing vertical jump ^(17)^. Simply extending the knees as rapidly as possible, with minimal force applied to the ground, is likely counterproductive for achieving optimal performance during a kettlebell swing. For older adults with painful arthritic knees, forcibly extending the joints may also exacerbate unwanted symptoms.

### Rate of force development

Data from proficient swings, shows a strong negative correlation (*r* = -0.82) in RFD with kettlebell mass ^(18)^, so it was surprising to find an almost neutral correlation among the novice older adults in this study (*r* = 0.03). The RFD effect size between the older adults in this study and instructor was large (δ = 1.47), with a mean difference of 17.0 N.s^-1^.kg^-1^; almost half that of the instructor (54.2%). With large within-group variation, it is interesting to consider the outliers. One male participant in the current study, who was still engaged in active employment, recorded a higher RFD than the instructor with the 12 kg kettlebell (δ = 0.25). This outlier shows that older adults can maintain a high RFD into older age and highlights the potential scope for others to improve. Results from strength and power training in older adults ^(63)^, indicate that it is feasible that kettlebell swings might increase RFD. Combined with large improvements in dynamic single leg balance ^(64)^ and postural reaction time ^(65)^, kettlebell training may mitigate age-related changes in RFD ^(66)^ and reduce falls risk. Whether swings might provide superior clinical benefit in this regard to a more basic exercise, such as the kettlebell deadlift, remains to be seen.

### Swing cadence

It was expected that swing cadence among novice participants would be lower than 40 SPM. There was a very large (δ = 2.54) average effect size between older adults and instructor ^(18)^, with the older adults cadence being on average 6 SPM slower. It was surprising that cadence was different between sexes - almost 2 SPM higher in females than males. Kettlebell-proficient females of comparable height, have previously been reported to perform swings at 40 SPM ^(67)^, thus anthropometric differences appear to have minimal influence on swing cadence. It was encouraging to see that cadence remained relatively consistent across all loads throughout the group, suggesting that the pre-selected kettlebells were not ‘too heavy’. In current practice, it is unlikely that insufficiently physically active females over 60 years of age would be asked to swing a 12-14 kg kettlebell, or men over 60 years asked to swing 16 kg. Findings from this exploratory investigation challenge common practice regarding kettlebell selection for older adults.

### Force time profile

The representative FTCs are more consistent with those of Mache and Hsieh ^(61)^ and Lake and Lauder ^(26)^ than of a proficient swing ^(18)^. Our observations support the findings of Back and colleagues ^(24)^ with one or both phases of ‘lift’ and ‘float’ typically being absent. Visual observation of the participant’s swing technique, and deviations from “ideal”, could be correlated with changes in the corresponding FTC. A common strategy for some of the participants in the present study, was to actively flex the shoulders to lift the kettlebell upward rather than swing it, which corresponded with a double-peak FTC. A second smaller force peak, seen in a proficient swing ^(18)^, was likely absent if the person was observed to control the kettlebell’s decent from mid-swing instead of allowing it to drop. Practically, this strategy shifts emphasis of the mechanical load of deceleration from the hips and lower limbs to the trunk and upper limbs.

Less visually apparent was the squatting motion previously observed in novices ^(24)^. This lack of squatting motion in the present study, was attributed to the instruction provided by the instructor and the pre-test drills which encouraged a ‘hip-hinge’ pattern. A wider, multi-peaked FTC and slower cadence was consistent with a more pronounced simultaneous, movement pattern of the lower limbs and trunk. Whether differences in swing pattern are clinically meaningful has not been investigated. Given the differences in force profile between a proficient and novice swing, it is hypothesised that that swing technique may influence performance improvement and rehabilitation conditions. An ‘ideal’ swing may not however be achievable for some older individuals, and variability in technique may also be beneficial.

As anticipated, none of the participants were able to perform an ideal hardstyle swing consistently, for all repetitions with all loads. There were, however, many examples where it was evident in the FTC, that participants had performed one or more swings within the set, which were approaching ideal – see Appendix 8: M4, M6, M7, M9, M12, M13, M15, M16, M17, F1, F3, F5, F9, F11-15, F17, F18. Most participants had a force profile suggestive of a swing rather than lift, which was surprising and very encouraging. Use of real-time biofeedback, in addition to common drills and slow-motion video analysis, may be beneficial for novices learning how to perform a hardstyle swing ^(68)^.

The hardstyle swing is an unusual movement pattern, which can be challenging for some individuals to learn, with older age likely to influence their movement patterns and potentially the rate of skill acquisition. Movement drills can be effective for improving technique, but some people can find them confusing; a situation which can be compounded by multiple drills. Many of the participants in the present study, however, were able to quickly improve their technique by simply focusing on increasing swing cadence. Swing cadence could be a useful measure to determine an appropriate upper limit for the most appropriate kettlebell, particularly in combination with an increasing rate of observed or reported effort. Achieving a consistent cadence of 40 SPM with a comfortable (lighter) kettlebell appears to be an appropriate strategy for the novice individual learning the swing. Health professionals should note that the cardiovascular demand of kettlebell swings may be higher than walking ^(37)^ and should anticipate a higher heart rate and cardiovascular response ^(69, 70)^ and program for the individual accordingly.

Our suggestion that changes in a FTC might be used to determine an optimal training load ^(18)^, assumes a proficient swing and availability of real time biofeedback. So how should optimal kettlebell load be determined for novice older adults? For some participants in the present study, increasing kettlebell mass appeared to improve their technique, but for others, additional mass had a detrimental effect. It remains to be seen whether monitoring the force profile to inform kettlebell selection provides any benefit beyond observation by a trained instructor. Optimising starting kettlebells, and progression during the initial and early stages of learning a hardstyle swing, is where input from an appropriately experienced coach is likely to be most beneficial.

The influence of confounding variables such as technique, age, and training history, on resultant force are unclear. Pragmatically however, the combined target testing loads per person for the deadlift session were 1,728 kg and 2,448 kg for females and males, respectively. With many older adults routinely under-dosed in resistance training programs ^(71)^, these data might help change that trend. In the absence of identifiable risks, and within the constraints of a person’s unique health history, clinical needs, and the desire (or not) to engage in a program of resistance training, these findings might encourage providers to broaden their perceived scope of therapeutic load. Additionally, clinical decision making is likely to influence instruction around movement, speed of movement, and treatment goals ^(42)^.

### Limitations

Results from insufficiently active older adults may not be generalisable to all older adults, or to those with a diagnosed health or medical condition which presents an increased risk of harm from resistance exercise, or which effects their ability to perform the exercise. Ground reaction force and technique among novices is likely to quickly improve with practice and appropriate instruction, therefore, these data may not represent older adults who have undergone a period of kettlebell exercise practice and training. Rate of force development should be interpreted with caution. Calculation of RFD from force-plate data, in the absence of concurrent video analysis, may have limited reliability, where determination of the propulsive phase in a force-time curve may not be clear. Ground reaction data from hardstyle swings cannot be generalised to the double knee-bend swing (kettlebell Sport) or overhead (American) swing which are kinematically different ^(36, 72)^.

## Conclusion

Results of this exploratory profiling study suggest that the prescription of exercise intensity (kg) for kettlebell swings and kettlebell deadlifts for older adults, should not be determined by sex. Ground reaction force was larger during two-handed hardstyle swings with an 8 kg kettlebell, than deadlifts performed with 32 kg. A target cadence of 40 SPM is likely to positively influence RFD and GRF in novices, but further research is required to establish whether such cadences are achievable for most older adults and how variations in cadence may influence aspects of hardstyle technique and the clinical outcomes of interest in older adults. All older females and males comfortably performed multiple deadlifts up to 24 kg and 32 kg, respectively, thus, healthcare providers are encouraged to explore the potential of their older patients/clients in this regard.

### Plain language summary

The magnitude of peak ground reaction force is considerably larger during a kettlebell swing than a kettlebell deadlift when the mass of the kettlebell is equal. Peak ground reaction force during a two-handed kettlebell swing with 8 kg is comparable to that of a 32 kg deadlift. Where improving the ability produce higher GRF is the clinical measure of interest, kettlebell swings with 8 kg may provide a similar benefit to deadlifts with heavier loads, and be a more cost-effective, convenient, and appealing option for older adults and healthcare providers. During kettlebell swings and kettlebell deadlifts, there is negligible difference in normalised peak ground reaction force between males and females. Findings suggest that older adults may have the potential, relative to bodyweight, to double the magnitude of peak ground reaction force and rate of force development during a kettlebell swing.

## Acknowledgements

The authors would like to acknowledge Mr Benjamin Hindle for his support in the calculation of rate of force development, and Mrs Evelyne Rathbone for her direction with statistical analysis.

## Funding

This study was supported by an Australian Government Research Training Program Scholarship and will contribute towards a Higher Degree by Research Degree (Doctor of Philosophy).

## Contributions

NM recruited the participants and conducted the study, curated, and analysed the data, interpreted the results, conducted the formal analysis, and wrote the original draft. JK, BS and WH supported with ongoing consultation. JK and WH reviewed and provided revisions to earlier versions of the manuscript. All authors read and approved the final manuscript.

## Ethics declarations

### Ethics approval and consent to participate

Not applicable

### Consent for publication

Not applicable

### Competing interests

Neil J. Meigh is a Physiotherapist and hardstyle kettlebell instructor, with an online presence as The Kettlebell Physio. Justin W.L. Keogh is an academic editor for PeerJ. BS and WH declare that they have no competing interests.

### Rights and permissions

Open Access This article is distributed under the terms of the Creative Commons Attribution 4.0 International License (http://creativecommons.org/licenses/by/4.0/), which permits unrestricted use, distribution, and reproduction in any medium, provided you give appropriate credit to the original author(s) and the source, provide a link to the Creative Commons license, and indicate if changes were made. The Creative Commons Public Domain Dedication waiver (http://creativecommons.org/publicdomain/zero/1.0/) applies to the data made available in this article, unless otherwise stated.

## Notes

### Competing Interest Statement

The primary author is a Physiotherapist and hardstyle kettlebell instructor, with an online presence as The Kettlebell Physio.

### Summary of Updates

Changes following peer-review: *Results unchanged* *Mechanical demands* replaced with *force profile* Study aims clarified Additional background information provided Pairwise comparisons made explicit Statistical analyses clarified (*exploratory* nature of the study reinforced) Explanation provided why all participants did not meet all conditions (making RM ANOVA inappropriate) Linear regression calculation added Description of the hardstyle swing added Swing cadence calculation added Numerous minor alterations to improve copy-text. Plain language summary added

