## Supplementary file A for "Force profile of the two-handed hardstyle kettlebell swing in novice older adults: an exploratory profile"

**Supplementary file A:** Force-time curves during two-handed hardstyle swings with a 12 kg kettlebell

**1. Males >60yrs**

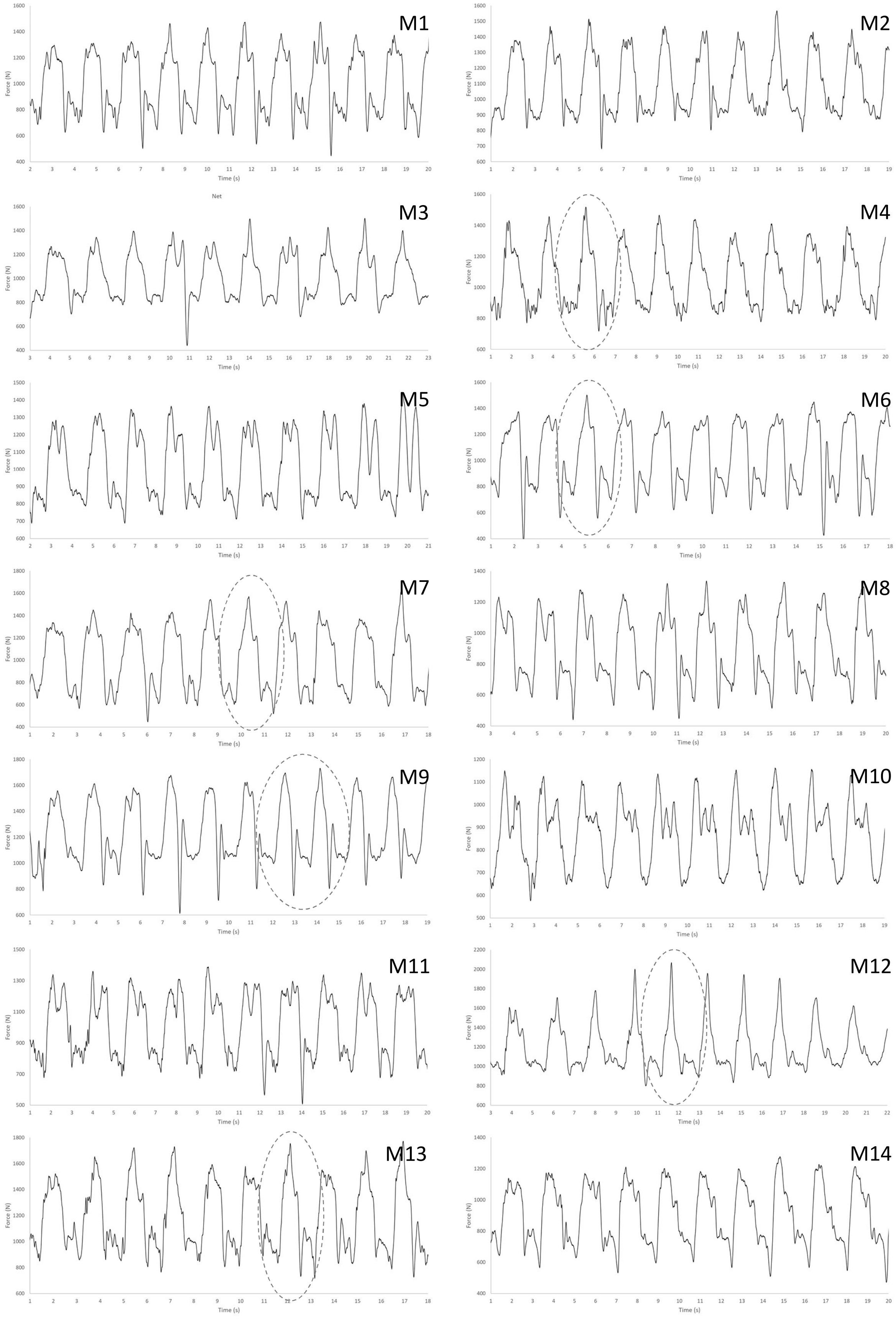

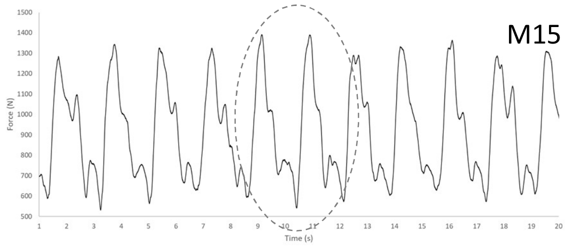
M15
Force (N)
Time (s)

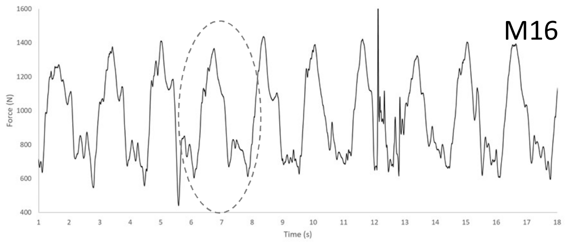
M16
Force (N)
Time (s)

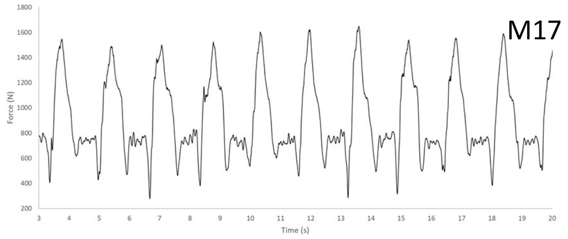
M17
Force (N)
Time (s)

**2. Females ≥59**

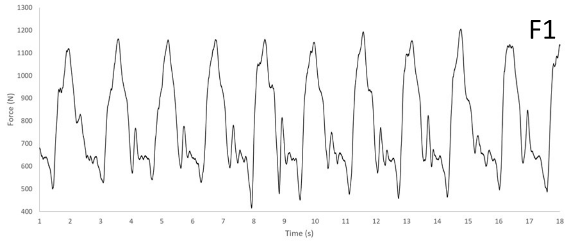
F1
Force (N)
Time (s)

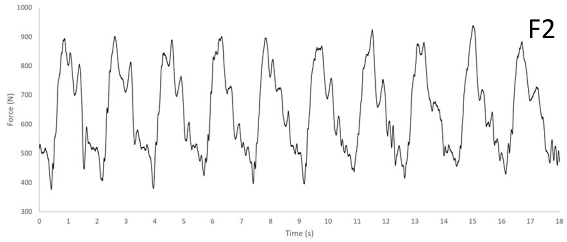
F2
Force (N)
Time (s)

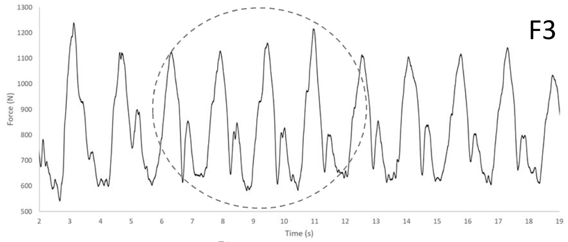
F3
Force (N)
Time (s)

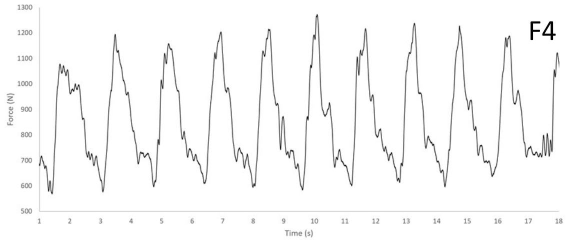
F4
Force (N)
Time (s)

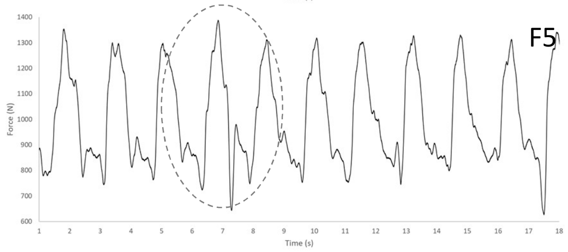
F5
Force (N)
Time (s)

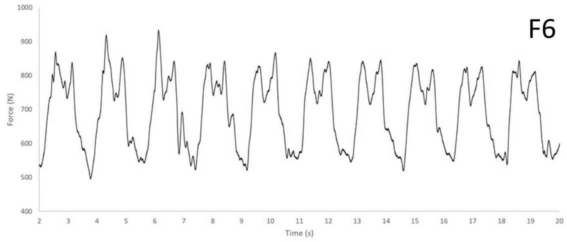
F6
Force (N)
Time (s)

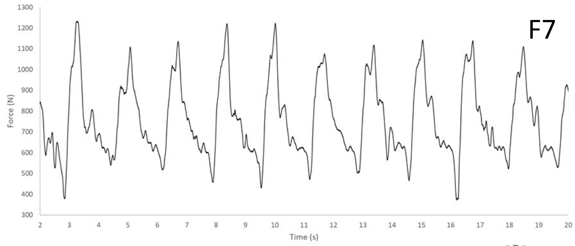
F7
Force (N)
Time (s)

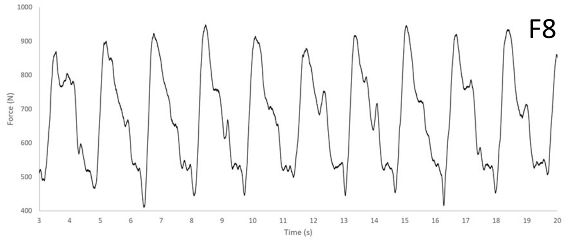
F8
Force (N)
Time (s)

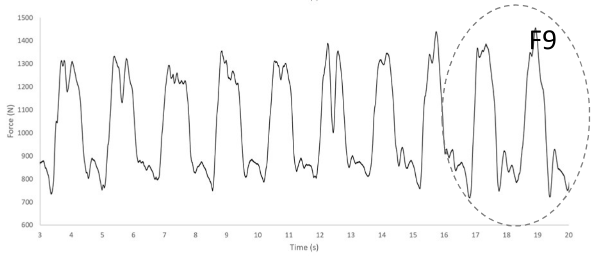
F9
Force (N)
Time (s)

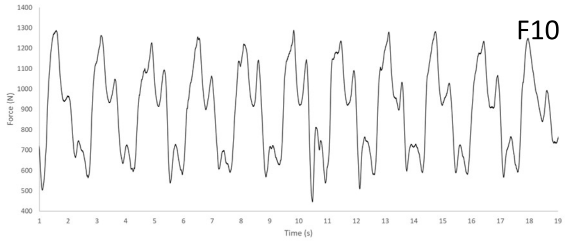
F10
Force (N)
Time (s)

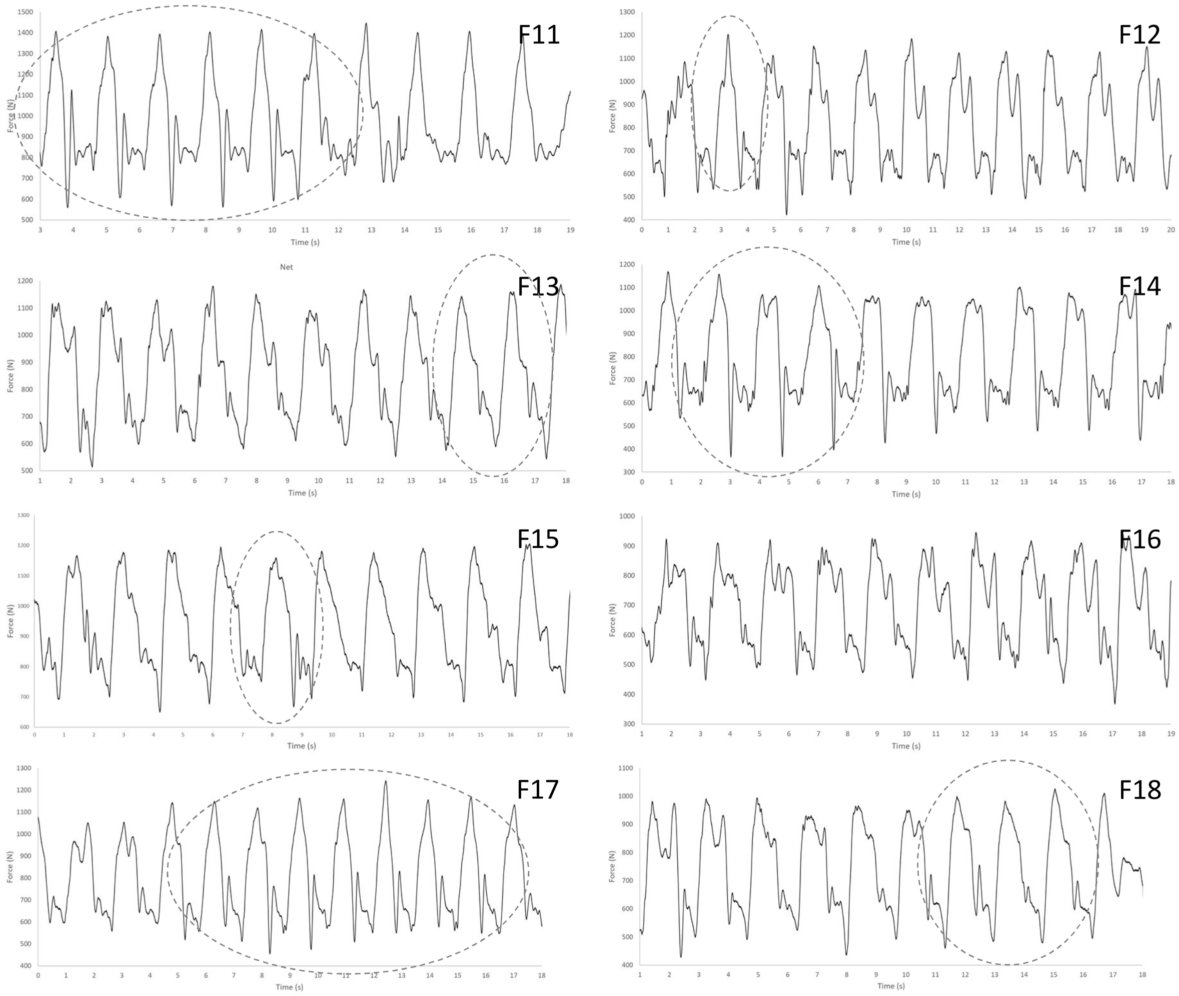
F11
Force (N)
Time (s)
Net
F12
Force (N)
Time (s)
F13
Force (N)
Time (s)
F14
Force (N)
Time (s)
F15
Force (N)
Time (s)
F16
Force (N)
Time (s)
F17
Force (N)
Time (s)
F18
Force (N)
Time (s)
